# Traits of soil bacteria predict plant responses to soil moisture

**DOI:** 10.1101/2022.04.08.487329

**Authors:** Lana G. Bolin, Jay T. Lennon, Jennifer A. Lau

## Abstract

Microbes can promote beneficial plant and animal responses to abiotic environments, but the ecological drivers of this benefit remain elusive. Here we investigated byproduct benefits, which occur when traits that increase the fitness of one species provide incidental benefits to another species with no direct cost to the provider species. In experimental mesocosms, microbial traits predicted plant responses to soil moisture such that bacteria with self-beneficial traits in drought increased plant early growth, size at reproduction, and chlorophyll concentration under drought, while bacteria with self-beneficial traits in well-watered environments increased these same plant traits in well-watered environments. Thus, microbial traits that promote microbial success in different soil moisture environments also promote plant success in these same environments. Our results show that the concept of byproduct benefits, originally conceived to explain the evolution of cooperation in pairwise mutualisms, also applies to interactions between plants and non-symbiotic soil microbes.

## INTRODUCTION

Plants and animals can adapt to a wide range of abiotic environments, but they often are not doing it alone. Instead, microbes can respond to abiotic environments in ways that help maintain plant and animal fitness via “microbial rescue” (*sensu* Mueller et al 2020; also called “microbe-mediated adaptation” *sensu* Petipas, Geber, and Lau 2021). For example, cold exposure can cause gut microbial community composition to shift in ways that promote cold tolerance in rodents (Chevalier et al. 2015), serpentine soil-adapted fungi can promote plant growth and phosphorus uptake in these phosphorus-limited soils (Doubková, Suda, and Sudová 2012), and endophytic fungi from hotter, drier environments can promote plant growth in drought (Giauque, Connor, and Hawkes 2019). The microbial benefits from these examples and others (*e*.*g*. Lau and Lennon 2012; Allsup and Lankau 2019; Fitzpatrick et al. 2018; Yuan et al. 2019) may help diverse plant and animal populations persist in stressful environments, but the ecological and evolutionary drivers of these patterns remain unresolved.

A strong foundation of theory informs understanding of the evolution of cooperation between closely interacting pairs of species. However, explaining the host fitness-promoting effects of non-symbiotic, diffusely interacting species is more challenging because it is unclear how the fitness of these species could become correlated (Sachs et al. 2004; Hawkes, Bull, and Lau 2020). One potential driver of microbial rescue between diffusely interacting species is byproduct benefits, which occur when the self-serving act of one species provides an incidental benefit to another species, with no direct cost to the provider species (Sachs et al. 2004).

Byproduct benefits can appear to be cooperative but are not in the classic sense because classic cooperation requires a cost to the provider species (Sachs et al. 2004). Other mechanisms that result in classic cooperation could also potentially contribute to beneficial interactions between diffuse, non-symbiotic partners. For example, partner choice could contribute if hosts can selectively reward the most beneficial non-symbiotic microbes, and partner fidelity feedbacks could contribute if hosts can selectively transfer beneficial non-symbiotic microbes to their offspring. However, classic cooperation may be less likely to occur when potential interactors re-assemble each generation (Hawkes, Bull, and Lau 2020; Sachs et al. 2004).

Byproduct benefits also may commonly promote microbial rescue because microbial communities are functionally diverse, allowing for rapid changes in microbial traits in different environments due either to rapid evolution or to changes in microbial community composition (Elena and Lenski 2003; Graves et al. 2015). The traits expressed in these changed microbial communities could then incidentally affect plant or animal responses to the environment. For example, herbicide application could select for microbes that degrade herbicide, which could then reduce soil herbicide stress for plants.

Here we investigated whether byproduct benefits can explain microbial rescue. We tested whether self-beneficial microbial traits under drought or well-watered conditions benefit plants in these same environments, and found evidence that byproduct benefits may explain previous observations of microbial communities responding to stress in ways that promote plant stress tolerance.

## METHODS

### Experimental Design Overview

We grew individuals of the annual legume *Chamaecrista fasciculata* (*Chamaecrista* hereafter) under well-watered or drought-stressed conditions and inoculated pots with one of 14 phylogenetically diverse bacterial strains representing three phyla and 12 families that varied in biofilm production and optimum water potential (N = 14 microbial strains × 2 watering treatments × 5 replicates = 180 plants; Appendix S1: Fig. S1, Fig. S2, Table S1). We then measured the influence of each bacterial trait on plant growth, as well as several plant traits that commonly respond to drought: leaf chlorophyll concentration, specific leaf area (SLA), timing of reproduction, and size at reproduction. Drought can reduce chlorophyll concentration by decreasing the lability of nutrients in dry soils (Evans 1989); early flowering is a common drought avoidance strategy; and lower SLA is associated with increased water use efficiency, making reduced SLA a putative adaptive response to drought (Ackerly 2004; Wright, Reich, and Westoby 2001).

The design ensured that any benefit conferred by microbes could derive only from byproduct benefits, and not classic cooperation. Classic cooperation requires repeated interactions between partners (“partner fidelity feedbacks”), multiple partners to choose from (“partner choice”), or close relatedness between partners (“kin selection”; Sachs et al. 2004). We eliminated these possibilities by using bacterial strains that are naïve to *Chamaecrista* (strains were isolated from bulk soil in an area where *Chamaecrista* did not occur) and by inoculating single strains, thereby preventing partner fidelity feedbacks and partner choice. If byproduct benefits occur, then we expect traits that are beneficial for microbes in drought and well-watered environments to benefit plants in those same environments.

### Bacterial Strains

The bacteria used in this experiment were isolated from bulk soils at the W.K. Kellogg Biological Station Long-Term Ecological Research site (KBS LTER, Hickory Corners, Michigan, USA). Strains were previously sequenced using the 16S rRNA gene and characterized for a range of functional traits, including biofilm production and optimum water potential (Lennon et al. 2012). Strains with low optimum water potentials achieve maximum growth in dry environments, making low optimum water potential an adaptive bacterial phenotype under drought, while high biofilm production is generally adaptive under drought because it reduces desiccation stress (Lennon et al. 2012; Lennon and Lehmkuhl 2016). Biofilm production was previously estimated using the Crystal Violet assay (O’Toole et al. 1999) and optimum water potential was estimated as the soil water potential at which a strain achieved maximum respiration rate (respiration rate strongly correlates with growth in these strains; see Lennon et al. 2012 for details). We selected 14 strains from this collection to maximize variation in biofilm production and optimum water potential. Therefore, although the traits correlated positively across the full collection of strains (Lennon et al. 2012), they were uncorrelated in our subset (*r* = −0.09, *P* = 0.64; Appendix S1: Fig. S1), allowing us to statistically partition the effects of each bacterial trait on plant traits. Prior to inoculation, each bacterial strain was grown in R2B medium (BD Difco, Sparks, Maryland, USA). Cells were then washed three times in phosphate-buffered saline. For details on strains and inoculum preparation, see Appendix S2.

### Greenhouse Experiment

We conducted a greenhouse experiment in fall 2019 in which we inoculated plants with each bacterial strain and grew them under drought or well-watered conditions. We sterilized and filled 656 mL Deepots™ (Stuewe and Sons, Tangent, Oregon, USA) with a sterile base soil composed of a 1:1 mixture of sand and our standard greenhouse mix (field soil mixed with organic material) that was twice steam sterilized (6 h at 77 °C with a 24 h rest between sterilizations), and we planted a single scarified and imbibed *Chamaecrista* seed (Prairie Moon Nursery, Winona, MN) into each pot. We then inoculated each bacterial strain onto ten replicate pots by pipetting 1 mL inoculum onto the thin layer of soil directly covering the seed. To aid in bacterial establishment, we reinoculated all pots 12 d after the initial inoculation (when most plants were at the three-leaf stage) by pipetting 1 mL suspension onto soil at the base of the plant.

We kept all plants well-watered for the first two weeks to promote plant establishment, at which point we began imposing drought stress on half of the replicates of each microbial inoculum treatment (n = 5 per strain). We watered all plants in the drought treatment only when they began showing signs of stress (*i*.*e*., wilting, 250 mL every ∼10 d), while we kept plants in the well-watered treatment well-watered throughout (250 mL approx. every ∼5 d). We fertilized all plants with a 1% solution 20-20-20 NPK fertilizer 7 and 10 weeks after planting.

### Plant Measurements

We measured six plant traits that commonly respond to drought: early growth, leaf chlorophyll concentration, specific leaf area (SLA), timing of reproduction, size at reproduction, and final biomass. Four weeks after planting, we counted the number of fully expanded leaves as an estimate of early growth, and measured leaf chlorophyll content as an indicator of plant nitrogen status (SPAD 502; Spectrum Technologies, Inc., Plainfield, IL, USA). Ten weeks after planting, we estimated SLA (leaf area/leaf dry weight) on the fifth fully expanded leaf on the main stem (or the nearest healthy leaf if the fifth was damaged) by measuring the area of each leaf on a leaf area meter (LI-3100C, Licor), then drying (60 °C for 14 d) and weighing. We recorded flowering date and the number of fully expanded leaves at the time of first flower as an estimate of plant size at reproduction. After all plants had flowered (10.5 w), we separated, dried (for at least 2 w at 60 °C), and weighed all shoot and root biomass.

### Statistical Analyses

To statistically control for microsite differences in the greenhouse, we created a detrended dataset by first regressing greenhouse block onto each plant trait, then adding each plant’s residual trait value to the global mean to return traits to their original scale.

To test whether bacterial biofilm production and optimum water potential predict plant responses to soil moisture, we fit phylogenetic generalized least squares (PGLS) models in R (nlme package v3.1-150; Garland et al. 1993; Martins and Hansen 1997; Pinheiro et al. 2018; R Core Team 2020). These models control for phylogenetic non-independence of the bacterial strains, but they cannot account for variation among individuals inoculated with a given strain (*i*.*e*., strain cannot be included as a random factor; we conducted additional analyses that account for variance within strains but that do not control for phylogenetic non-independence and detected weaker effects; Appendix S1: Table S2). Therefore, because the unit of replication is the bacterial strain, we conducted our analyses on strain means within each watering treatment (Huang, Lankau, and Peng 2018). Strains did not differ in variance for any plant traits except time of reproduction and final biomass, which were not significantly predicted by bacterial traits (see Results; Bartlett’s K^2^: all *P* > 0.3 except time of reproduction *P* = 0.047, and final biomass *P* = 0.002), suggesting that this was a reasonable approach. We constructed a phylogenetic tree using the 16S rRNA gene sequences of our strains generated by Lennon et al. (2012). We aligned these sequences (MUSCLE v3.8.31; Edgar 2004) and generated a maximum likelihood tree with 100 bootstrap replicates (RAxML v8.2.12; Stamatakis 2014) using the General Time Reversible (GTR) model of nucleotide substitution with a gamma distributed substitution rate. We then built the correlation structure of our tree that would be expected if the traits evolve under Brownian motion (ape package v5.4-1; Paradis and Schliep 2019) and fit PGLS models assuming this correlation structure.

We analyzed each plant response variable separately in models that included bacterial biofilm production, bacterial optimum water potential, watering treatment (drought or well-watered), and all interactions as fixed effects. We *ln*-transformed optimum water potential because water potential is a nonlinear function of volumetric water content such that small reductions in water content in dry soils are associated with large reductions in water potential (Bilskie and Scientific 2001). To do this we multiplied all values by −1 to make them positive (water potentials are always negative), took the natural logarithm of the positive value, then multiplied again by −1 to return values to their original order. We standardized both bacterial traits to a mean of zero and a variance of one to make coefficients comparable and to prevent the generation of spurious correlations between each trait and their interaction, which can happen when predictors are expressed on different scales (Aiken, West, and Reno 1991). We assessed statistical significance using Type III ANOVA (car package v3.1-10; Fox and Weisberg 2019).

## RESULTS

Bacterial traits predicted the magnitude of plant responses to soil moisture. Bacteria with low optimum water potentials (dry-adapted bacteria) reduced the negative effects of drought on plant early growth and size at reproduction, while bacteria with high optimum water potentials (wet-adapted bacteria) increased the benefits plants received from growing in well-watered conditions (drought × optimum, early growth: *χ*^*2*^_*1,20*_ = 8.3, *P* < 0.01; size at reproduction: *χ*^*2*^_*1,20*_ = 7.6, *P* < 0.01; Fig. 1A,B; Appendix S1: Table S3). As a result, drought reduced predicted plant early growth by 44% and size at reproduction by 49% when pots were inoculated with microbes with the highest optimum water potential, compared to only 32% and 30%, respectively, when inoculated with microbes with the lowest optimum water potential. Additionally, bacteria with the lowest optimum water potential resulted in smaller predicted declines in plant SLA caused by drought relative to bacteria with the highest optimum water potentials (drought × optimum: *χ*^*2*^_*1,20*_ = 4.3, *P* = 0.04; Fig. 1F, Appendix S1: Table S3).

**Figure 1.**
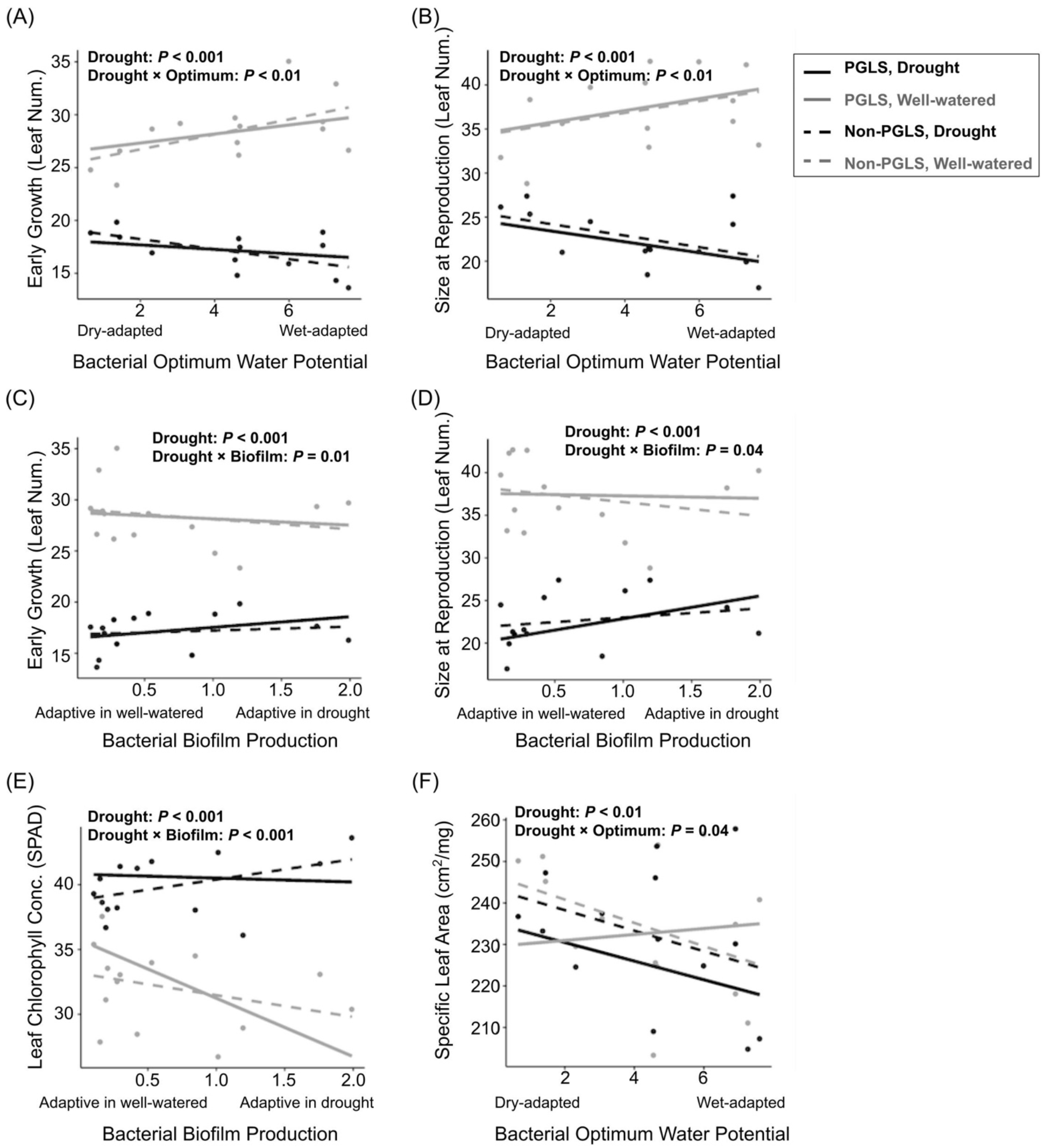
Bacterial optimum water potential (“Optimum”) predicted plant responses to soil moisture in (A) early growth, (B) size at reproduction, and (F) specific leaf area (SLA), while bacterial biofilm production (“Biofilm”) predicted plant responses to soil moisture in (C) early growth, (D), size at reproduction, and (E) chlorophyll concentration. Solid lines represent fitted phylogenetic least squares (PGLS) regressions, and dashed lines represent fitted regressions that do not control for bacterial phylogeny. Biofilm production is reported as the relative absorbance generated from the biofilm assay (Lennon et al. 2012). High biofilm production is adaptive for bacteria in drought, while low biofilm production is adaptive in well-watered environments. Optimum water potential was ln-transformed as described in the Methods. Bacteria with low optimum water potentials have high growth rates in drought, while bacteria with high optimum water potentials have high growth rates in well-watered environments. Reported *P*-values are from PGLS models.

Bacteria that produced large amounts of biofilm also mitigated the negative effects of drought on plants. Drought reduced predicted plant early growth by 42% and size at reproduction by 45% when pots were inoculated with microbes with the lowest biofilm production, compared to only 33% and 31%, respectively, when inoculated with microbes with the highest biofilm production (drought × biofilm, early growth: *χ*^*2*^_*1,20*_ = 6.2, *P* = 0.01; size at reproduction: *χ*^*2*^_*1,20*_ = 4.4, *P* = 0.04; Fig. 1C,D). Bacterial biofilm production also influenced plant chlorophyll responses to soil moisture (drought × biofilm: *χ*^*2*^_*1,20*_ = 30.1, *P* < 0.001; Fig. 1E; Appendix S1: Table S3). Specifically, well-watered plants had lower chlorophyll concentrations (drought: *χ*^*2*^_*1,20*_ = 243, *P* < 0.001), especially when inoculated with high biofilm-producing bacteria. This occurred because high biofilm-producing bacteria reduced chlorophyll in well-watered conditions (biofilm: *χ*^*2*^_*1,10*_ = 7.8, *P* < 0.01), while biofilm did not affect chlorophyll under drought (*χ*^*2*^_*1,10*_ = 0.03, *P* = 0.86). Note, however, that well-watered plants had more chlorophyll on a per plant basis (estimated as chlorophyll concentration per leaf × leaf number) than drought-stressed plants because they were larger (drought: *P* < 0.001; Fig. 1A; Appendix S1: Fig. S3).

The effect of each bacterial trait on plant chlorophyll and SLA responses to soil moisture depended on the other bacterial trait (drought × optimum × biofilm, SLA: *χ*^*2*^_*1,20*_ = 17.75, *P* < 0.001; Chlorophyll: *χ*^*2*^_*1,20*_ = 18.76,17.75, *P* < 0.001; Appendix S1: Fig. S4). However, these results should be interpreted cautiously because our data are not evenly distributed across the response surface (Albert et al. 2010).

Drought stress reduced plant final biomass by 36% (drought: *χ*^*2*^_*1,20*_ = 76.5, *P* < 0.001), and accelerated plant flowering by 4.5 days (drought: *χ*^*2*^_*1,20*_ = 9.1, *P* < 0.01; Appendix S1: Table S3), but these plant traits were not affected by bacterial traits.

## DISCUSSION

We showed that self-beneficial bacterial traits under drought (low optimum water potential and high biofilm production) also benefited plants under drought, while self-beneficial bacterial traits in well-watered soils (high optimum water potential and low biofilm production) also benefited plants in well-watered soils. These findings indicate that byproduct benefits can contribute to microbial rescue. This resulted in drought-stressed plants more closely resembling the well-watered phenotype in terms of plant growth when grown with bacteria with drought-ameliorating traits (Fig. 1A-D). These findings provide proof of concept that theory on the evolution of cooperation can explain fitness correlations between diffusely interacting pairs of species, and show how byproduct benefits can be detected using experimental systems that eliminate the possibility of classic cooperation.

Our results highlight the potential importance of microbial traits in plant responses to the environment. In fact, additional comparisons to sterile inocula revealed that trait effects were stronger than the effects of live vs. sterile inoculation, which did not predict any plant responses to soil moisture (Appendix S3: Fig. S1, Table S1). This suggests that studies manipulating live vs. sterile inoculation may be missing important microbial effects on plants that are mediated by microbial traits.

Additionally, our results provide strong support for the utility of a predictive trait-based approach. While ecological models are reasonably good at predicting how traits will respond to different environments (*e*.*g*., Grime 1977; Chapin 1980; Litchman et al. 2007; Pedley and Dolman 2014), and to a lesser extent how traits can affect other organisms in the community (*e*.*g*., Baxter et al. 2019), we have shown that microbial response traits measured in a lab can predict not only the traits of other community members, but also how those community members’ traits respond to the environment.

We focused on microbial rescue in response to soil moisture, but there is no reason to expect that this is a soil moisture-specific phenomenon. A similar pattern could emerge in any environment where microbial traits that are adaptive for microbes also benefit plants. For example, herbicide application has been shown to select for microbes that degrade herbicides (El Fantroussi, Verschuere, and Verstraete 1999; Lancaster et al. 2010), and this microbial trait could benefit plants by removing herbicides from the soil. Such plant benefit mediated by diffuse communities of microbes may be widespread but is rarely investigated.

### Microbial traits predict plant soil moisture responses: potential mechanisms

Byproduct benefits from microbial traits might promote beneficial plant responses to soil moisture by at least three non-mutually exclusive mechanisms: microbial trait expression could directly manipulate plant phenotype (*e*.*g*., production of phytohormones; Yang, Kloepper, and Ryu 2009), modify the soil environment (*e*.*g*., alter soil water holding capacity; Lennon and Lehmkuhl 2016; Martiny et al. 2015), or allow generally beneficial microbes to survive and maintain their beneficial function (*e*.*g*., help rhizobia survive and continue fixing nitrogen for plants in exchange for carbon). We did not attempt to differentiate among mechanisms in our study, but we speculate that biofilm production may have increased plant early growth and size at reproduction under drought by increasing soil water holding capacity (Lennon and Lehmkuhl 2016). Increased survival of generally beneficial microbes seems less likely, as inoculation did not increase any plant growth metrics relative to uninoculated pots (Appendix S1: Fig. S5), and while we cannot rule out plant hormone-mediated effects, we have no *a priori* reason to predict that bacterial phytohormone production would correlate with biofilm production.

The mechanism underlying the impact of microbes with low optimum water potentials on plant drought responses is less clear. These bacteria were not generally beneficial to plants, but it is unclear how other traits known to contribute to this moisture niche besides biofilm production, such as dormancy (Lennon and Jones 2011), solute production (Schimel, Balser, and Wallenstein 2007), and the ability to change intracellular stoichiometry (Fredrickson, Shu-mei, and Gaidamakova 2008), would increase plant growth or SLA, or alter phytohormone production.

Two of our strains belong to genera with known plant growth-promoting properties: a *Burkholderia* species and a *Pseudomonas* species (Hayat, Ahmed, and Sheirdil 2012). However, these strains did not drive microbial rescue in either watering treatment. While these strains often increased plant growth relative to predicted growth for a given biofilm production or optimum water potential, particularly under drought, they almost always weakened microbial trait effects (Appendix S1: Fig. S6). These findings support our conclusions that the traits of these microbes, and not their identities, drove microbial rescue in response to soil moisture.

For microbial traits to benefit plants, one might expect microbial traits would shift plant traits in an adaptive manner. Thicker leaves (low SLA) are putatively adaptive under drought (Ackerly 2004; Wright, Reich, and Westoby 2001), and we found that plants produced thicker leaves under drought. However, the magnitude of effect was greatest when grown with microbes with high, not low, optimum water potentials. Rather than altering the expression of plant traits in an adaptive manner, perhaps the protective benefits of these microbes simply reduced the need for plants to plastically produce thicker leaves in response to drought. Indeed, microbes with low optimum water potentials generally caused drought-stressed plants to resemble well-watered plants, suggesting a protective effect (Fig. 1A-D).

### Caveats –

We tested for byproduct benefits using a simple system of single strain inoculations on a single plant species. We did this to better isolate the effects of microbial traits, but it is unclear whether these effects will scale up to more complex communities found in nature. Whole microbial communities certainly vary in biofilm production and other traits that may benefit plants under drought (*e*.*g*., Berard et al. 2015), but microbial trait expression is complex and could be strongly affected by other microbial community members (Classen et al. 2015). In fact, a small pilot study involving these same strains did not find consistent effects of simple communities comprised of high vs. low biofilm producing strains on plant drought responses (Appendix S4: Fig. S1).

We showed that byproduct benefits can promote microbial rescue, but it remains to be seen how commonly this occurs in nature, how these effects on individual performance scale up to populations, and how important microbial rescue is relative to other forms of drought response such as plasticity or adaptive evolution. Even within microbial rescue, modes of classic cooperation (*e*.*g*., partner choice and partner fidelity feedbacks) could contribute alongside byproduct benefits. Quantifying the relative importance of these drivers could be a fruitful area for future research.

### Implications & Conclusions

If microbial traits commonly influence plant responses to the abiotic environment, and if our results scale up to diverse microbial communities, then microbial rescue driven by byproduct benefits could be common. Because microbial traits can respond rapidly to changes in the environment, sometimes in as little as a single week (*e*.*g*., Mackelprang et al. 2011), microbial traits could promote plant population persistence under a range of short-term events like seasonal drought, as well as facilitate plant persistence under longer-term global changes (Hawkes, Bull, and Lau 2020; Petipas, Geber, and Lau 2021). Ultimately, our results show that byproduct benefits are a feasible mechanism that could promote commonly observed, but previously challenging to explain, patterns of microbial rescue in both plants and animals.

## ACKNOWLEDGMENTS

LGB was supported as an NSF Graduate Research Fellow. We thank Brent Lehmkuhl for lab assistance and consultation in microbial inocula preparation; Nicole Hardy, Lauren Martin, and Kai Smith for help collecting data; John Lemon and Tom Pirtle for greenhouse assistance; and Jennifer Rudgers and the Lau lab for thoughtful comments on this manuscript. This work was funded in part by the NSF Long-Term Ecological Research Program at the Kellogg Biological Station (NSF DEB 1832042), the National Science Foundation (DEB-1934554 JTL, DBI-2022049 JTL, BCS-2009125 JAL and JTL), a US Army Research Office Grant (W911NF-14-1-0411 JTL), the National Aeronautics and Space Administration (80NSSC20K0618 JTL), and by Michigan State University AgBioResearch. This is KBS contribution #2311.

## LITERATURE CITED

Ackerly, David. 2004. “Functional Strategies of Chapparal Shrubs in Relation to Seasonal Water Deficit and Disturbance.” Ecological Monographs 74 (1): 25–44.

Aiken, Leona S., Stephen G. West, and Raymond R. Reno. 1991. Multiple Regression: Testing and Interpreting Interactions. SAGE.

Albert Cécile H., Nigel G. Yoccoz, Thomas C. Edwards Jr, Catherine H. Graham, Niklaus E. Zimmermann, and Wilfried Thuiller. 2010. “Sampling in Ecology and Evolution - Bridging the Gap between Theory and Practice.” Ecography 33 (6): 1028–37.

Allsup, Cassandra, and Richard Lankau. 2019. “Migration of Soil Microbes May Promote Tree Seedling Tolerance to Drying Conditions.” Ecology 100 (9): e02729.

Baxter, Nielson T., Alexander W. Schmidt, Arvind Venkataraman, Kwi S. Kim, Clive Waldron, and Thomas M. Schmidt. 2019. “Dynamics of Human Gut Microbiota and Short-Chain Fatty Acids in Response to Dietary Interventions with Three Fermentable Fibers.” mBio 10 (1).

Berard, Annette, Meriem Ben Sassi, Aurore Kaisermann, and Pierre Renault. 2015. “Soil Microbial Community Responses to Heat Wave Components: Drought and High Temperature.” Climate Research 66 (3). https://doi.org/10.3354/cr01343.

Bilskie, Jim, and Campbell Scientific. 2001. “Soil Water Status: Content and Potential.” Campbell Scientific, Inc. App. Note: 2S–1.

Chapin, F. S., III. 1980. “The Mineral Nutrition of Wild Plants.” Annual Review of Ecology and Systematics 11 (1): 233–60.

Chevalier, Claire, Ozren Stojanović, Didier J. Colin, Nicolas Suarez-Zamorano, Valentina Tarallo, Christelle Veyrat-Durebex, Dorothée Rigo, et al. 2015. “Gut Microbiota Orchestrates Energy Homeostasis during Cold.” Cell 163 (6): 1360–74.

Classen Aimée T., Maja K. Sundqvist, Jeremiah A. Henning, Gregory S. Newman, Jessica A. M. Moore, Melissa A. Cregger, Leigh C. Moorhead, and Courtney M. Patterson. 2015. “Direct and Indirect Effects of Climate Change on Soil Microbial and Soil Microbial-Plant Interactions: What Lies Ahead?” Ecosphere 6 (8): art130.

Doubková, Pavla, Jan Suda, and Radka Sudová. 2012. “The Symbiosis with Arbuscular Mycorrhizal Fungi Contributes to Plant Tolerance to Serpentine Edaphic Stress.” Soil Biology & Biochemistry 44 (1): 56–64.

Edgar, Robert C. 2004. “MUSCLE: Multiple Sequence Alignment with High Accuracy and High Throughput.” Nucleic Acids Research 32 (5): 1792–97.

Elena, Santiago F., and Richard E. Lenski. 2003. “Microbial Genetics: Evolution Experiments with Microorganisms: The Dynamics and Genetic Bases of Adaptation.” Nature Reviews. Genetics 4 (6): 457.

El Fantroussi, S., L. Verschuere, and W. Verstraete. 1999. “Effect of Phenylurea Herbicides on Soil Microbial Communities Estimated by Analysis of 16S rRNA Gene Fingerprints and Community-Level Physiological Profiles.” Appl. Environ.

Evans, John R. 1989. “Photosynthesis and Nitrogen Relationships in Leaves of C3 Plants.” Oecologia 78 (1): 9–19.

Fitzpatrick, Connor R., Julia Copeland, Pauline W. Wang, David S. Guttman, Peter M. Kotanen, and Marc T. J. Johnson. 2018. “Assembly and Ecological Function of the Root Microbiome across Angiosperm Plant Species.” Proceedings of the National Academy of Sciences of the United States of America 115 (6): E1157–65.

Fox, John, and Sanford Weisberg. 2019. “An R Companion to Applied Regression (Third).” Thousand Oaks CA: Sage.

Fredrickson, J. K., W. L. Shu-mei, and E. K. Gaidamakova. 2008. “Protein Oxidation: Key to Bacterial Desiccation Resistance?” The ISME Journal.

Garland, Theodore, Jr, Allan W. Dickerman, Christine M. Janis, and Jason A. Jones. 1993. “Phylogenetic Analysis of Covariance by Computer Simulation.” Systematic Biology 42 (3): 265–92.

Giauque, Hannah, Elise W. Connor, and Christine V. Hawkes. 2019. “Endophyte Traits Relevant to Stress Tolerance, Resource Use and Habitat of Origin Predict Effects on Host Plants.” The New Phytologist 221 (4): 2239–49.

Graves, Joseph L., Jr, Mehrdad Tajkarimi, Quincy Cunningham, Adero Campbell, Herve Nonga, Scott H. Harrison, and Jeffrey E. Barrick. 2015. “Rapid Evolution of Silver Nanoparticle Resistance in Escherichia Coli.” Frontiers in Genetics 6 (February): 42.

Grime, J. P. 1977. “Evidence for the Existence of Three Primary Strategies in Plants and Its Relevance to Ecological and Evolutionary Theory.” The American Naturalist 111 (982): 1169–94.

Hawkes, Christine V., James J. Bull, and Jennifer A. Lau. 2020. “Symbiosis and Stress: How Plant Microbiomes Affect Host Evolution.” Philosophical Transactions of the Royal Society of London. Series B, Biological Sciences 375 (1808): 20190590.

Hayat, Rifat, Iftikhar Ahmed, and Rizwan Ali Sheirdil. 2012. “An Overview of Plant Growth Promoting Rhizobacteria (PGPR) for Sustainable Agriculture.” In Crop Production for Agricultural Improvement, edited by Muhammad Ashraf, Münir Öztürk, Muhammad Sajid Aqeel Ahmad, and Ahmet Aksoy, 557–79. Dordrecht: Springer Netherlands.

Huang, Fangfang, Richard Lankau, and Shaolin Peng. 2018. “Coexistence via Coevolution Driven by Reduced Allelochemical Effects and Increased Tolerance to Competition between Invasive and Native Plants.” The New Phytologist 218 (1): 357–69.

Lancaster, Sarah H., Emily B. Hollister, Scott A. Senseman, and Terry J. Gentry. 2010. “Effects of Repeated Glyphosate Applications on Soil Microbial Community Composition and the Mineralization of Glyphosate.” Pest Management Science 66 (1): 59–64.

Lau, Jennifer A., and Jay T. Lennon. 2012. “Rapid Responses of Soil Microorganisms Improve Plant Fitness in Novel Environments.” Proceedings of the National Academy of Sciences of the United States of America 109 (35): 14058–62.

Lennon, Jay T., Zachary T. Aanderud, B. K. Lehmkuhl, and Donald R. Schoolmaster Jr. 2012. “Mapping the Niche Space of Soil Microorganisms Using Taxonomy and Traits.” Ecology 93 (8): 1867–79.

Lennon, Jay T., and Stuart E. Jones. 2011. “Microbial Seed Banks: The Ecological and Evolutionary Implications of Dormancy.” Nature Reviews. Microbiology 9 (2): 119–30.

Lennon, Jay T., and Brent K. Lehmkuhl. 2016. “A Trait-Based Approach to Bacterial Biofilms in Soil.” Environmental Microbiology 18 (8): 2732–42.

Litchman, Elena, Christopher A. Klausmeier, Oscar M. Schofield, and Paul G. Falkowski. 2007. “The Role of Functional Traits and Trade-Offs in Structuring Phytoplankton Communities: Scaling from Cellular to Ecosystem Level.” Ecology Letters 10 (12): 1170–81.

Mackelprang, Rachel, Mark P. Waldrop, Kristen M. DeAngelis, Maude M. David, Krystle L. Chavarria, Steven J. Blazewicz, Edward M. Rubin, and Janet K. Jansson. 2011. “Metagenomic Analysis of a Permafrost Microbial Community Reveals a Rapid Response to Thaw.” Nature 480 (7377): 368–71.

Martins, Emilia P., and Thomas F. Hansen. 1997. “Phylogenies and the Comparative Method: A General Approach to Incorporating Phylogenetic Information into the Analysis of Interspecific Data.” The American Naturalist 149 (4): 646–67.

Martiny, Jennifer B. H., Stuart E. Jones, Jay T. Lennon, and Adam C. Martiny. 2015. “Microbiomes in Light of Traits: A Phylogenetic Perspective.” Science 350 (6261).

Mueller, Emmi A., Nathan I. Wisnoski, Ariane L. Peralta, and Jay T. Lennon. 2020. “Microbial Rescue Effects: How Microbiomes Can Save Hosts from Extinction.” Functional Ecology 34 (10): 2055–64.

O’Toole George A., Leslie A. Pratt, Paula I. Watnick, Dianne K. Newman, Valerie B. Weaver, and Roberto Kolter. 1999. “[6] Genetic Approaches to Study of Biofilms.” In Methods in Enzymology, 310:91–109. Academic Press.

Paradis, Emmanuel, and Klaus Schliep. 2019. “Ape 5.0: An Environment for Modern Phylogenetics and Evolutionary Analyses in R.” Bioinformatics 35 (3): 526–28.

Pedley, Scott M., and Paul M. Dolman. 2014. “Multi-Taxa Trait and Functional Responses to Physical Disturbance.” The Journal of Animal Ecology 83 (6): 1542–52.

Petipas, Renee H., Monica A. Geber, and Jennifer A. Lau. 2021. “Microbe-Mediated Adaptation in Plants.” Ecology Letters 24 (7): 1302–17.

Pinheiro, J., D. Bates, S. DebRoy, and D. Sarkar. 2018. “R Core Team. 2018. Nlme: Linear and Nonlinear Mixed Effects Models. R Package Version 3.1-137.” URL: https://CRAN.R-Project.Org/package=Nlme.

R Core Team (2020). R: A language and environment for statistical computing. R Foundation for Statistical Computing, Vienna, Austria. URL https://www.R-project.org/.

Sachs, Joel L., Ulrich G. Mueller, Thomas P. Wilcox, and James J. Bull. 2004. “The Evolution of Cooperation.” The Quarterly Review of Biology 79 (2): 135–60.

Schimel, Joshua, Teri C. Balser, and Matthew Wallenstein. 2007. “Microbial Stress-Response Physiology and Its Implications for Ecosystem Function.” Ecology 88 (6): 1386–94.

Stamatakis, Alexandros. 2014. “RAxML Version 8: A Tool for Phylogenetic Analysis and Post-Analysis of Large Phylogenies.” Bioinformatics 30 (9): 1312–13.

Wright, I. J., P. B. Reich, and M. Westoby. 2001. “Strategy Shifts in Leaf Physiology, Structure and Nutrient Content between Species of High- and Low-Rainfall and High- and Low-Nutrient Habitats.” Functional Ecology 15 (4): 423–34.

Yang, Jungwook, Joseph W. Kloepper, and Choong-Min Ryu. 2009. “Rhizosphere Bacteria Help Plants Tolerate Abiotic Stress.” Trends in Plant Science 14 (1): 1–4.

Yuan, Yongge, Caroline Brunel, Mark van Kleunen, Junmin Li, and Zexin Jin. 2019. “Salinity-Induced Changes in the Rhizosphere Microbiome Improve Salt Tolerance of Hibiscus Hamabo.” Plant and Soil 443 (1): 525–37.

