## Supplementary Material for "Traits of soil bacteria predict plant responses to soil moisture"

Microbial traits explain microbe-mediated host adaptation

**Appendix S1**

**Figure S1.** Biofilm production and optimum water potential are uncorrelated in the 14 bacterial strains used in our study. Each point represents a bacterial strain. Optimum water potential was ln-transformed by multiplying all values by −1 to make them positive (water potentials are always negative values), taking the natural logarithm of the positive value, then multiplying again by −1 to return values to their original order.

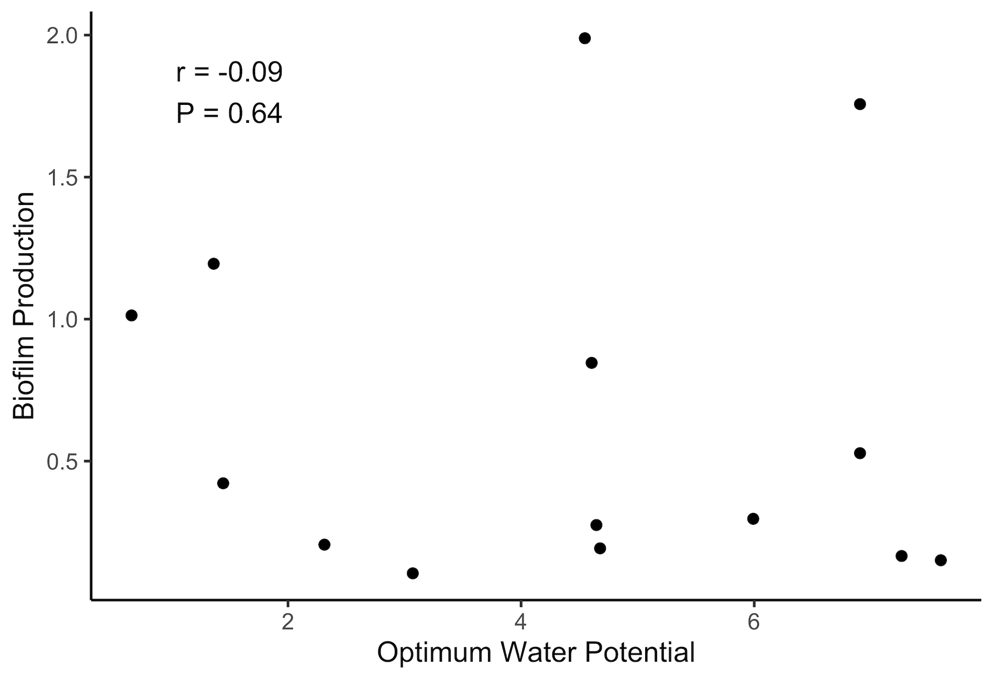

**Figure S2.** Phylogenetic tree showing the phylogenetic and trait diversity of bacterial strains used in this study. Tree was constructed from aligned 16S rRNA sequences using maximum likelihood (RAxML) methods. Numbers associated with nodes are ML bootstrap frequencies. Shaded boxes represent standardized bacterial trait values (biofilm production and −(ln(−optimum water potential))). Lines to the right of the branches indicate strains that fall within a given phylum.

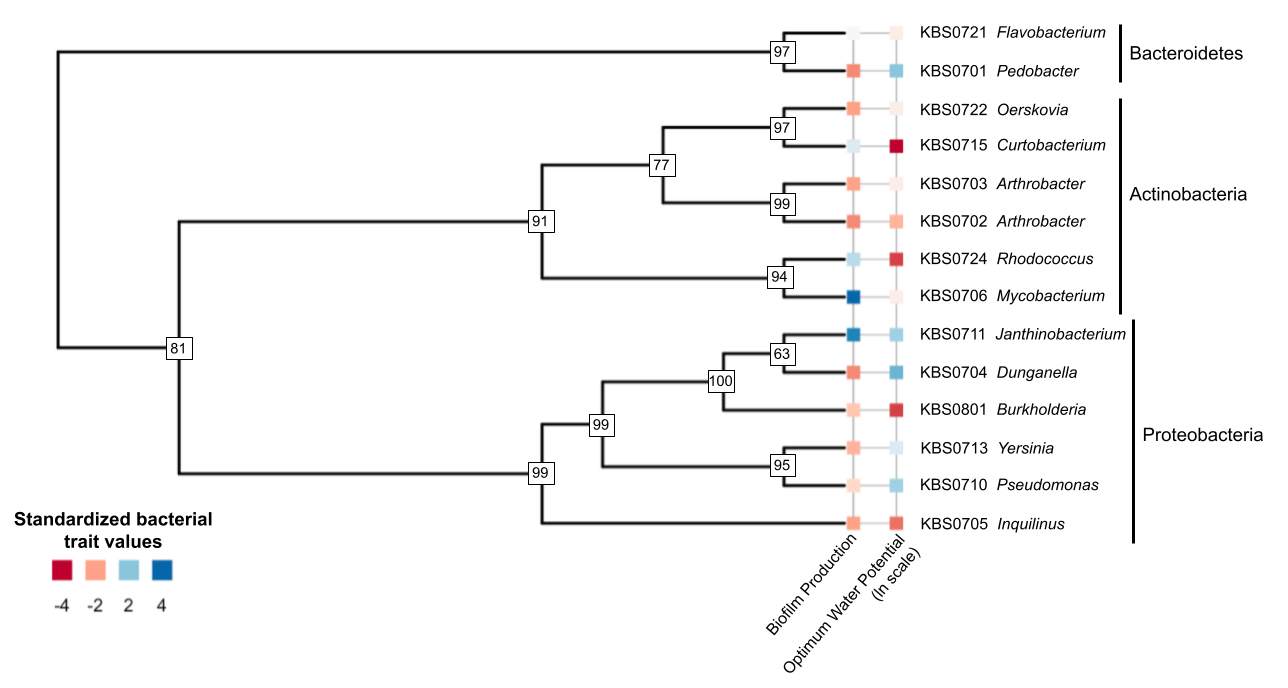

**Figure S3.** Estimates of per-plant chlorophyll (calculated by multiplying chlorophyll concentration per leaf by the number of leaves) were greater in wet soil than in dry soil. Solid lines represent fitted phylogenetic least squares (PGLS) regressions, and dashed lines represent fitted regressions that do not control for bacterial phylogeny. Biofilm production is reported as the relative absorbance generated from the biofilm assay (Lennon et al. 2012).

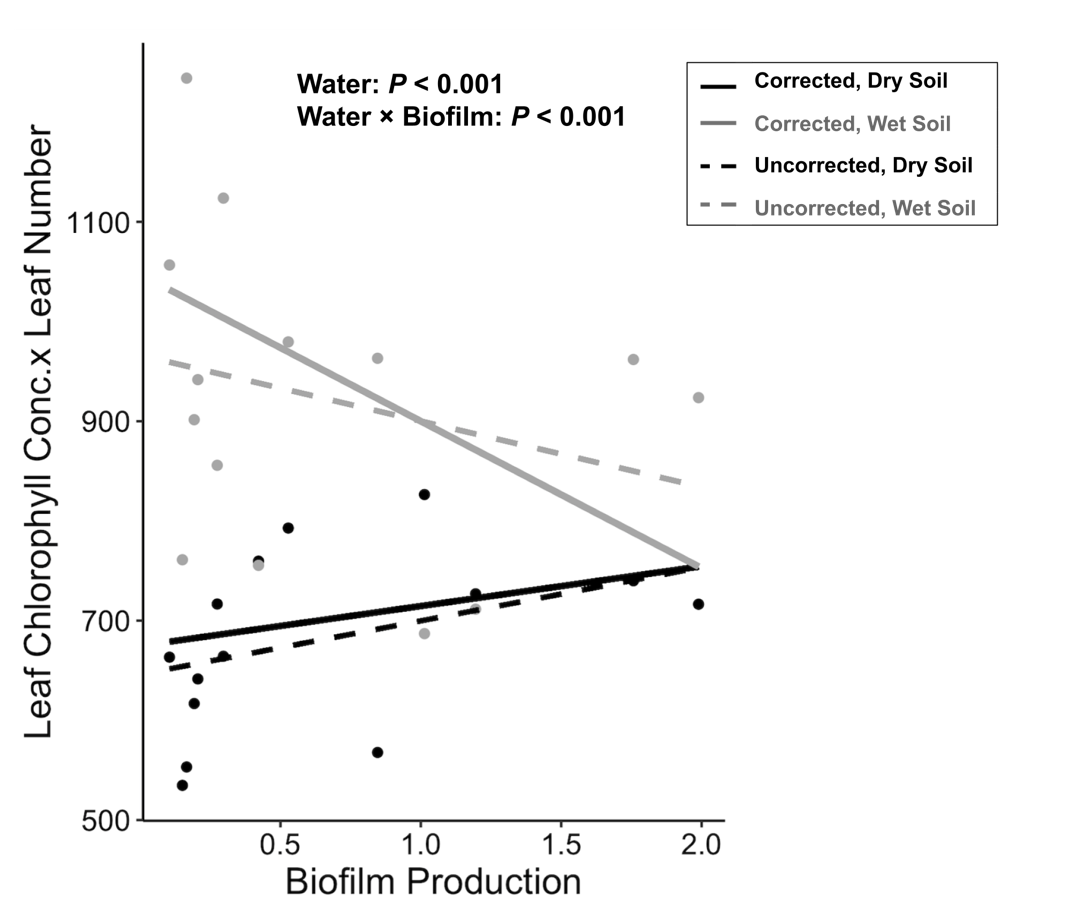

**Figure S4.** 3D plots showing the significant water × optimum water potential × biofilm production interaction for leaf chlorophyll concentration (top row) and specific leaf area (bottom row) in dry (left column) and wet (right column) conditions.

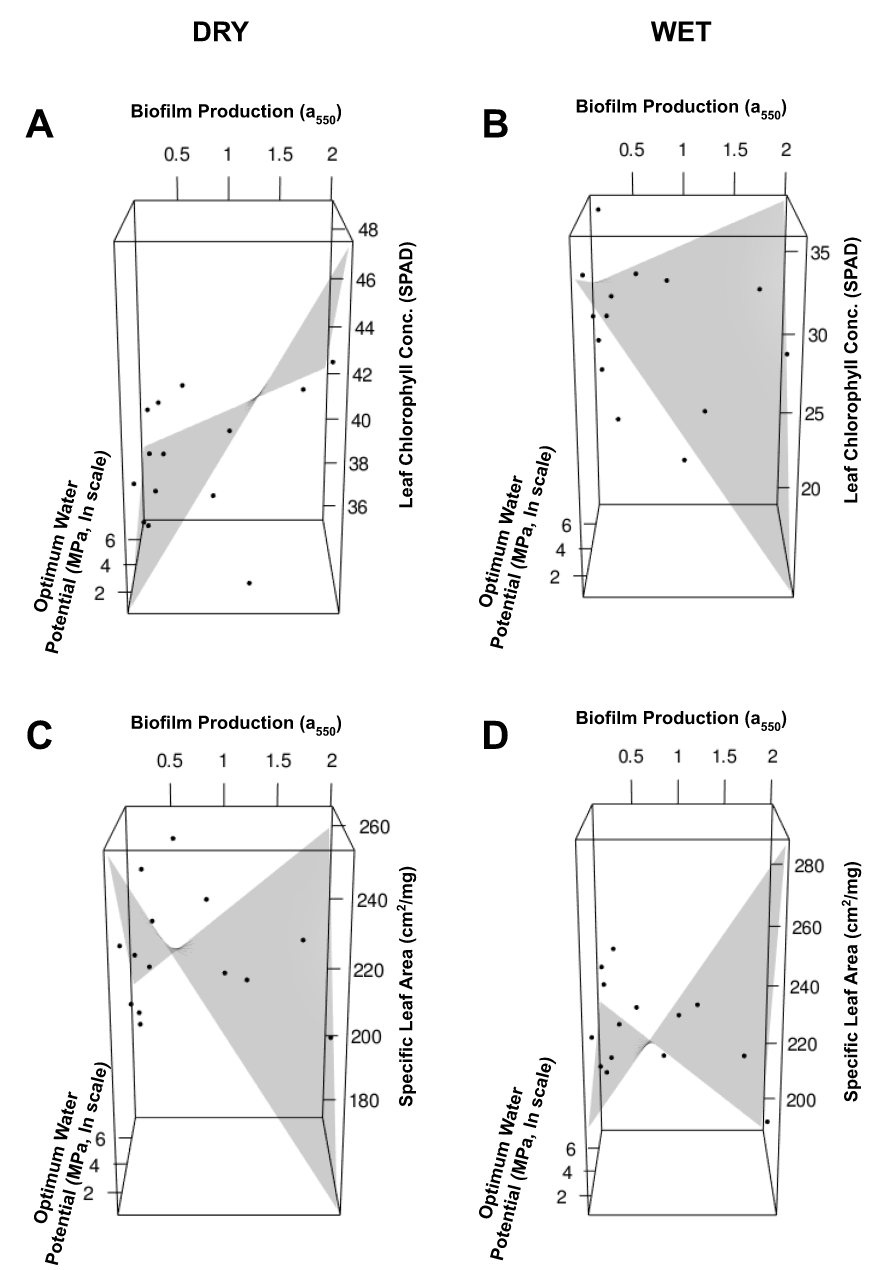

**Figure S5.** No strains provided a significant growth benefit over controls in terms of plant (a) early growth, (b) size at reproduction, or (c) final biomass (t-test: all *P* > 0.1). Controls are shown on the far left in red, followed by the 14 bacterial strains used in our study. Error bars represent SE.

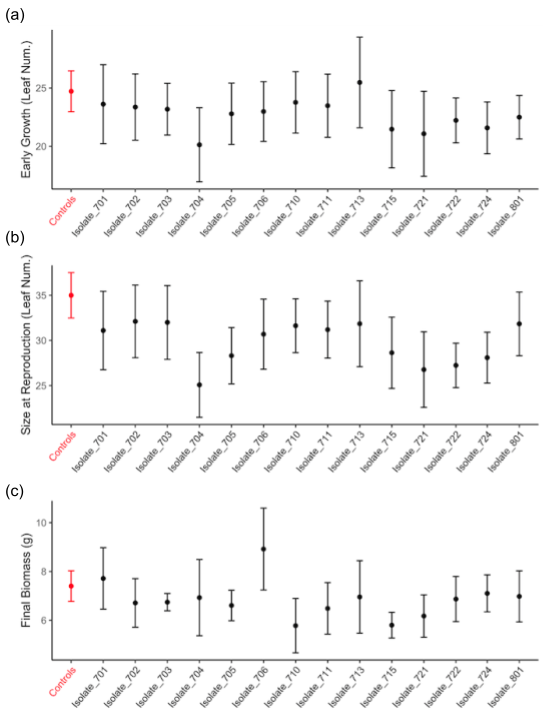

**Figure S6.** Plant responses to soil moisture were predicted by bacterial traits. This figure is identical to Figure 1 in the main text, except strains belonging to genera with known plant growth-promoting properties – a *Burkholderia* species and a *Pseudomonas* species – are labeled. Burk. = *Burkholderia*, Pseud. = *Pseudomonas*.

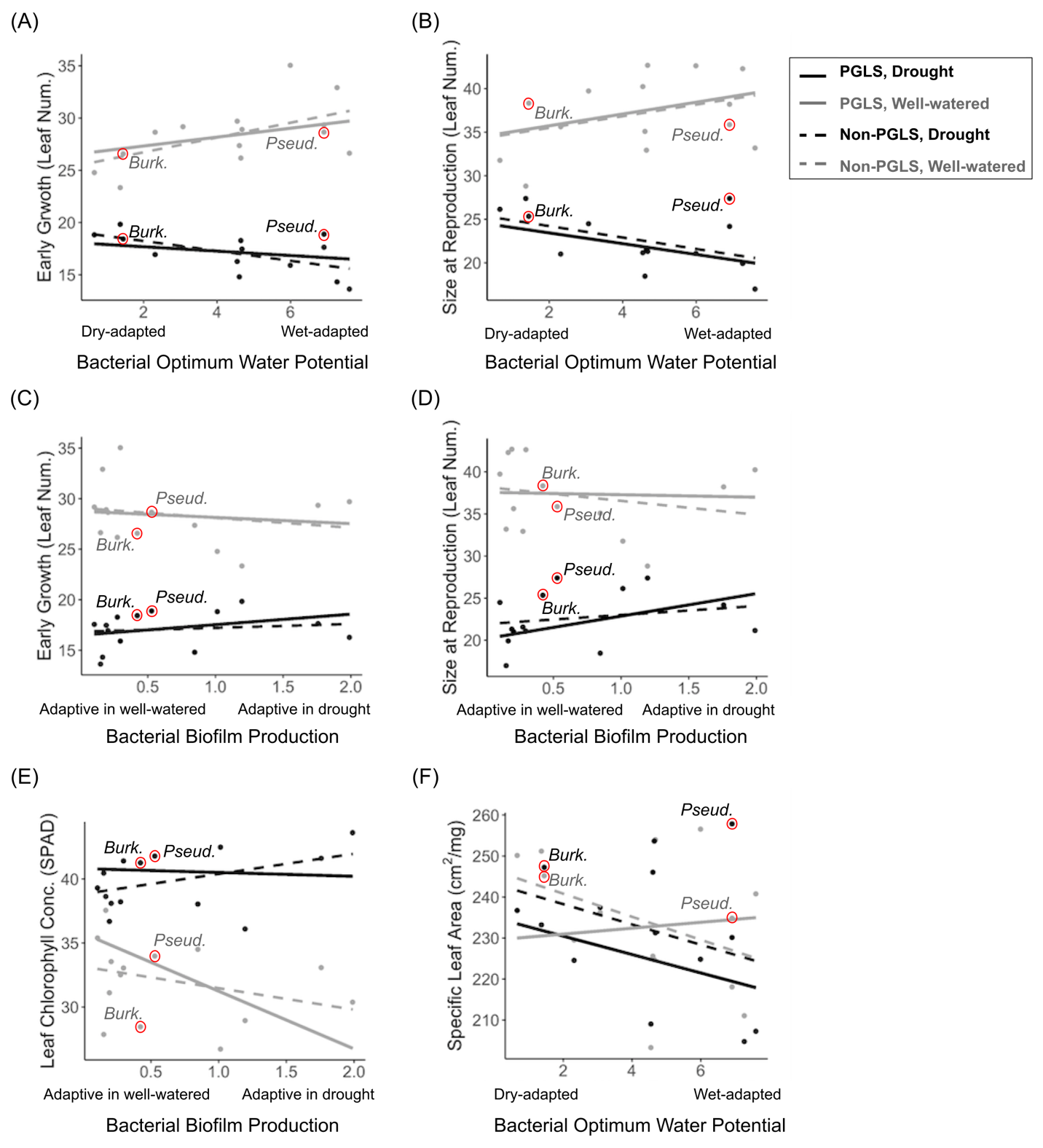

**Table S1.** Phylogenetic and trait information for each bacterial isolate. KBS Code is the unique identifier for each strain and is consistent with previous studies using these strains. Subsequent columns list taxonomic information for each isolate, followed by the biofilm production and optimum water potential of each isolate as measured by Lennon et al. (2012)**.** Accession number can be used to retrieve isolate DNA sequences from GenBank ([www.ncbi.nlm.nih.gov](http://www.ncbi.nlm.nih.gov)). Adapted from Lennon et al. (2012).

| **KBS Code** | **Domain** | **Phylum** | **Class** | **Order** | **Family** | **Genus** | **Biofilm Production (a_550_)** | **Optimum Water Potential (-MPa)** |
| --- | --- | --- | --- | --- | --- | --- | --- | --- |
| 0701 | Bacteria | Bacteroidetes | Sphingobacteria | Sphingobacteriales | Sphingobacteriaceae | *Pedobacter* | 0.166 | -0.0007 |
| 0702 |  | Actinobacteria | Actinobacteria | Actinomycetales | Micrococcaceae | *Arthrobacter* | 0.105 | -0.0464 |
| 0703 |  | Actinobacteria | Actinobacteria | Actinomycetales | Micrococcaceae | *Arthrobacter* | 0.193 | -0.0093 |
| 0704 |  | Proteobacteria | β-Proteobacteria | Burkholderiales | Oxalobacteraceae | *Dunganella* | 0.151 | -0.0005 |
| 0705 |  | Proteobacteria | α-Proteobacteria | Rhodospirillales | Rhodospirillaceae | *Inquilinus* | 0.206 | -0.0990 |
| 0706 |  | Actinobacteria | Actinobacteria | Actinomycetales | Mycobacteriaceae | *Mycobacterium* | 1.989 | -0.0106 |
| 0710 |  | Proteobacteria | γ-Proteobacteria | Pseudomonadales | Pseudomonadaceae | *Pseudomonas* | 0.528 | -0.0010 |
| 0711 |  | Proteobacteria | β-Proteobacteria | Burkholderiales | Oxalobacteraceae | *Janthinobacterium* | 1.757 | -0.0010 |
| 0713 |  | Proteobacteria | γ-Proteobacteria | Enterobacteriales | Enterobacteriaceae | *Yersinia* | 0.297 | -0.0025 |
| 0715 |  | Actinobacteria | Actinobacteria | Actinomycetales | Microbacteriaceae | *Curtobacterium* | 1.013 | -0.5180 |
| 0721 |  | Bacteroidetes | Flavobacteria | Flavobacteriales | Flavobacteriaceae | *Flavobacterium* | 0.846 | -0.0100 |
| 0722 |  | Actinobacteria | Actinobacteria | Actinomycetales | Cellulomonadaceae | *Oerskovia* | 0.275 | -0.0096 |
| 0724 |  | Actinobacteria | Actinobacteria | Actinomycetales | Nocardiaceae | *Rhodococcus* | 1.195 | -0.2560 |
| 0801 |  | Actinobacteria | β-Proteobacteria | Burkholderiales | Burkholderiaceae | *Burkholderia* | 0.422 | -0.2360 |

**Table S2.** Results from linear mixed models (lme4; Bates et al. 2015) that account for variance within strains, but that do not control for phylogenetic non-independence. Models included watering treatment (“water”: drought-stressed or well-watered), bacterial biofilm production (“biofilm”), bacterial optimum water potential (“optimum”), and all interactions as fixed effects, and bacterial strain as a random effect. *** *P* < 0.001, ** *P* < 0.01, * *P* < 0.05, † *P* < 0.1.

|  | **df** | **Early Growth**  **𝝌^2^** | **Size at Reproduction**  **𝝌^2^** | **Chlorophyll Conc.**  **𝝌^2^** | **Days to First Flower**  **𝝌^2^** | **SLA**  **𝝌^2^** | **Biomass**  **𝝌^2^** |
| --- | --- | --- | --- | --- | --- | --- | --- |
| **Intercept** | 1 | 432 *** | 423 *** | 2510*** | 1.16e12 *** | 2150 *** | 195 *** |
| **Water** | 1 | 98.0 *** | 91.9 *** | 43.5*** | 1.66 | 0.10 | 28.3 *** |
| **Biofilm** | 1 | 0.07 | 0.35 | 1.41 | 0.04 | 0.52 | 0.001 |
| **Optimum** | 1 | 1.77 | 1.88 | 0.25 | 0.001 | 1.63 | 0.22 |
| **Water × Biofilm** | 1 | 0.47 | 1.07 | 2.89 † | 0.37 | 0.12 | 1.18 |
| **Water × Optimum** | 1 | 4.29 * | 2.45 | 0.11 | 1.60 | 0.06 | 1.01 |
| **Biofilm × Optimum** | 1 | 0.10 | 0.03 | 0.03 | 0.41 | 0.88 | 0.08 |
| **Water × Biofilm × Optimum** | 1 | 0.13 | 0.84 | 0.84 | 0.03 | 1.37 | 0.06 |
| **Residual** | 20 |  |  |  |  |  |  |

**Table S3.** Results from phylogenetic generalized least squares (PGLS) models testing the effects of watering treatment (“water”: drought-stressed or well-watered), bacterial biofilm production (“biofilm”), and bacterial optimum water potential (“optimum”) on plant traits. *** *P* < 0.001, ** *P* < 0.01, * *P* < 0.05, † *P* < 0.1.

|  | **df** | **Early Growth**  **𝝌^2^** | **Size at Reproduction**  **𝝌^2^** | **Chlorophyll Conc.**  **𝝌^2^** | **Days to First Flower**  **𝝌^2^** | **SLA**  **𝝌^2^** | **Biomass**  **𝝌^2^** |
| --- | --- | --- | --- | --- | --- | --- | --- |
| **Intercept** | 1 | 0.25 | 0.09 | 1.05 | 0.70 | 1.03 | 0.04 |
| **Water** | 1 | 667*** | 281*** | 243*** | 9.13** | 7.52** | 76.5*** |
| **Biofilm** | 1 | 0.003 | 0.004 | 0.0002 | 0.004 | 0.002 | 0.001 |
| **Optimum** | 1 | 0.003† | 0.005 | 0.003 | 0 | 0.006 | 0.001 |
| **Water × Biofilm** | 1 | 6.24* | 4.43* | 30.1*** | 0.08 | 0.73 | 2.06 |
| **Water × Optimum** | 1 | 8.31** | 7.62** | 0.91 | 1.28 | 4.29* | 0.07 |
| **Biofilm × Optimum** | 1 | 0.001 | 0.002 | 0.02 | 0.005 | 0 | 0.0001 |
| **Water × Biofilm × Optimum** | 1 | 2.28 | 0.02 | 18.8*** | 3.63† | 17.75*** | 0.005 |
| **Residual** | 20 |  |  |  |  |  |  |

**Appendix S2**

BACTERIAL STRAINS AND INOCULUM PREP

The 14 bacterial taxa used in this experiment represent diverse strains belonging to 11 families, 13 genera, and 3 phyla (Figure S2; Table S1). They were isolated from the KBS LTER in 2007 and 2008, and have since been cryogenically preserved in 20% glycerol at −80°C.

To prepare single-strain inocula, we revived each of the cryogenically preserved isolates by plating them onto R2 agar. We then inoculated a single colony of each isolate into 50 ml of R2B liquid medium which we incubated on a shaker table at 25 °C for 24 h. We washed each liquid culture by pelleting the cells three times in phosphate-buffered saline (PBS), then resuspended each culture in 50 ml of PBS before inoculating plants. We additionally prepared four sterilized control inocula to test how inoculation with live bacteria influences plant drought responses. One control was microbe-free PBS. For the other three controls, we autoclaved additional inocula from three of the isolates (the lowest and highest biofilm production [0.105 and 1.989 a_550_, respectively], as well as an intermediate biofilm-producing isolate [0.528 a_550_]) at 121 °C for 20 min before inoculating plants.

**Appendix S3**

LIVE INOCULATION ANALYSES AND RESULTS

To test whether inoculation with live bacteria predicts plant traits in dry and wet environments, we fit a linear model for each plant trait that included sterilization treatment (live or sterilized inoculum), watering treatment (dry or wet) and their interaction as fixed effects. These analyses did not correct for phylogeny because

Plants that were inoculated with one of the four control inocula (sterilized medium biofilm-producing isolate) behaved significantly differently than the other controls (plant early growth and size at reproduction were higher when inoculated with this control in wet conditions relative to the other controls; water × inoculum type: early growth *F* = 3.6, *P* = 0.03; size at reproduction *F* = 2.9, *P* = 0.05). Therefore, we present results from the more conservative models that exclude this control.

***Inoculation with live bacteria did not predict plant responses to soil moisture*** (sterilization treatment × water: all *P* > 0.1; Table S1). However, live inoculation caused plants to flower 3.0 days earlier (sterilization treatment: *F* = 7, *P* = 0.01; Fig. S1d) and at a 11.5% smaller size (sterilization treatment: *F =* 4.9, *P* = 0.03; Fig. S1b; Table S1) relative to plants inoculated with sterile controls. Sterilization treatment did not influence any other plant traits.

**Figure S1.** Live inoculation did not influence plant responses to soil moisture (water × sterilization: all *P* > 0.18). Live inoculation (d) accelerated plant flowering and (b) caused plants to flower at a smaller size relative to plants inoculated with sterilized controls, but did not affect any other plant traits (a,c,e,f). Error bars are fitted SE.

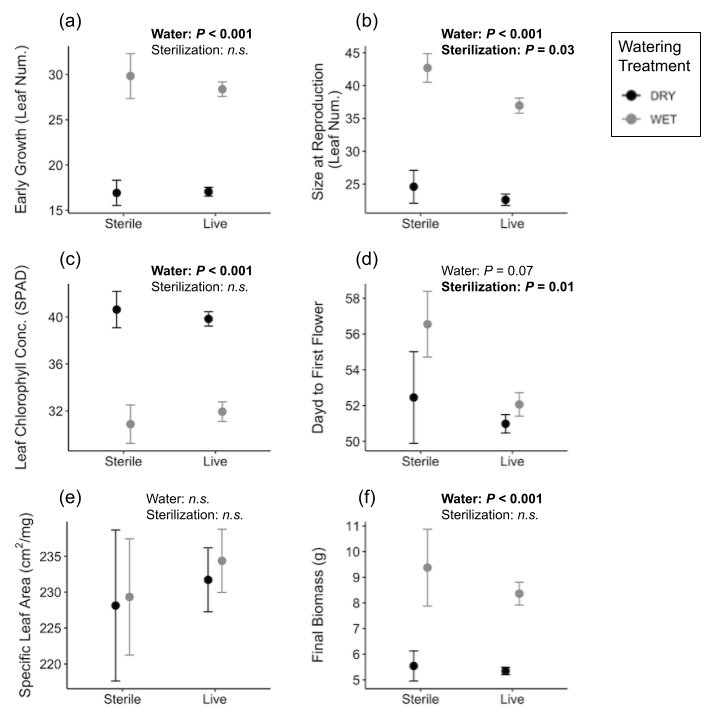

**Table S1.** F-statistics and associated *P*-values from general linear models testing the effects of watering treatment (“water”: drought-stressed or well-watered) and sterilization treatment (live or sterile inoculation) on plant traits. *** *P* < 0.001, ** *P* < 0.01, * *P* < 0.05, † *P* < 0.1.

|  | **df** | **Early Growth**  **F** | **Size at Reproduction**  **F** | **Chlorophyll Conc.**  **F** | **Days to First Flower**  **F** | **SLA**  **F** | **Biomass**  **F** |
| --- | --- | --- | --- | --- | --- | --- | --- |
| **Water** | 1 | 162*** | 128*** | 76.4*** | 3.55† | 0.18 | 45.9*** |
| **Sterilization Treatment** | 1 | 0.30 | 4.93* | 0.01 | 7.04* | 0.33 | 0.98 |
| **Water × Sterilization Treatment** | 1 | 0.44 | 1.15 | 0.57 | 1.80 | 0.01 | 0.45 |
| **Residual** | 30 |  |  |  |  |  |  |

**Appendix S4**

COMMUNITY BIOFILM PILOT STUDY RESULTS

We grew *Chamaecrista* individuals under well-watered or drought-stressed conditions, and inoculated them with three types of simple, four-strain bacterial communities (N = 3 community types × 20 replicates × 2 watering treatments = 120 plants). For community type A, we randomly selected four strains from the six lowest biofilm producing strains from our collection. For community type B, we randomly selected four strains from the six highest biofilm producing strains. For community type C, we randomly selected two strains from the six lowest biofilm producing strains and two strains from the six highest biofilm producing strains. We prepared inocula as described in Appendix B, except community inocula were composed of four strains in equal proportion instead of a single strain. The rest of the experiment was conducted identically to the experiment described in the main text.

To test whether biofilm production in simple bacterial communities predicts plant responses to soil moisture, we fit linear mixed models in R. We analyzed the means of each unique inoculum to account for non-independence of each four-strain community, for a total of 10 data points per community per watering treatment. We analyzed each plant trait separately in models that included mean biofilm production (calculated as the mean biofilm production of the four individual strains in each inoculum), watering treatment (dry or wet), and their interaction as fixed effects, and we included greenhouse block as a random effect. We assessed statistical significance using Type III ANOVA with Satterthwaite’s approximation of denominator degrees of freedom (car package v3.1-10; Fox and Weisberg 2019).

Mean community biofilm production influenced plant size at reproduction responses to soil moisture, and tended to influence plant early growth responses to soil moisture (water × biofilm, size at reproduction: *F_1,56_* = 6.2, *P* = 0.02; early growth: *F_1, 56_* = 2.8, *P* = 0.098; Fig. S1a,b). Greater mean biofilm production tended to increase plant size at reproduction in wet soils (mean biofilm: *F_1,28_* = 3.9, *P* = 0.06), which is opposite the effect seen with single strain inocula. However, mean biofilm production did not influence plant size at reproduction in dry soils. Mean biofilm production did not significantly influence plant early growth in either soil moisture environment.

Mean biofilm production also influenced plant flowering time responses to soil moisture (water × biofilm: *F_1,56_* = 3.8, *P* = 0.047; Fig. S1c), although it did not significantly influence flowering time in either soil moisture environment. These results, too, are in contrast to those of single strain inocula where biofilm production did not influence plant flowering time.

Mean biofilm production did not influence plant chlorophyll concentration, contrary to results with single strain inocula (water × biofilm: *F_1,56_* = 0.11, *P* = 0.74).

**Figure S1.**

Mean bacterial biofilm production and watering treatment effects on plan (a) early growth, (b) size at reproduction, and (c) days to first flower. Biofilm bacterial community biofilm production is reported as the mean relative absorbance of the four strains included in each inoculum (generated from the biofilm assay; Lennon et al. 2012).

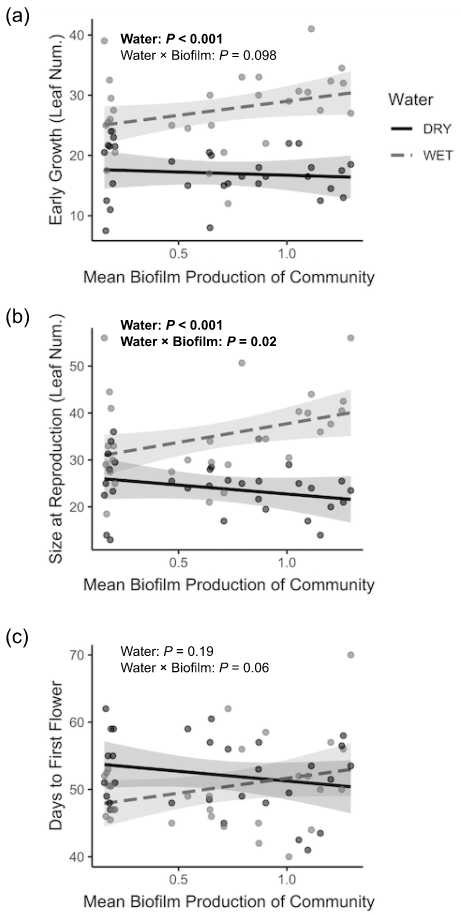

**LITERATURE CITED**

Bates, Douglas, Martin Maechler, Ben Bolker, Steven Walker, Rune Haubo Bojesen Christensen, Henrik Singmann, Bin Dai, Gabor Grothendieck, Peter Green, and Maintainer Ben Bolker. 2015. “Package ‘lme4.’” *Convergence* 12 (1): 2.

Lennon, Jay T., Zachary T. Aanderud, B. K. Lehmkuhl, and Donald R. Schoolmaster Jr. 2012. “Mapping the Niche Space of Soil Microorganisms Using Taxonomy and Traits.” *Ecology* 93 (8): 1867–79.
